# Prenatal opioid exposure inhibits microglial sculpting of the dopamine system selectively in adolescent male offspring

**DOI:** 10.1101/2021.11.28.468091

**Authors:** Caroline J. Smith, Tania Lintz, Madeline J. Clark, Karen E. Malacon, Nicholas J. Constantino, Veronica J. Kim, Young C. Jo, Yanaira Alonso-Caraballo, Alia Abiad, Staci D. Bilbo, Elena H. Chartoff

**Affiliations:** Department of Psychiatry, Harvard Medical School and Basic Neuroscience Division, Mclean Hospital, Belmont, MA; Department of Psychology and Neuroscience, Duke University, Durham, NC

**Keywords:** Microglia, dopamine, phagocytosis, prenatal opioid, oxycodone, nucleus accumbens

## Abstract

The current opioid epidemic has dramatically increased the number of children who are prenatally exposed to opioids, including oxycodone. A number of social and cognitive abnormalities have been documented in these children as they reach young adulthood. However, little is known about the mechanisms underlying developmental effects of prenatal opioid exposure. Microglia, the resident immune cells of the brain, respond to acute opioid exposure in adulthood. Moreover, microglia are known to sculpt neural circuits during healthy development. Indeed, we recently found that microglial phagocytosis of dopamine D1 receptors (D1R) in the nucleus accumbens (NAc) is required for the natural developmental decline in NAc-D1R that occurs between adolescence and adulthood in rats. This microglial pruning occurs only in males, and is required for the normal developmental trajectory of social play behavior. However, virtually nothing is known as to whether this developmental program is altered by prenatal exposure to opioids. Here, we show in rats that maternal oxycodone self-administration during pregnancy leads to reduced adolescent microglial phagocytosis of D1R and subsequently higher D1R density within the NAc in adult male, but not female, offspring. Finally, we show that prenatal opioid exposure abolishes the extinction of oxycodone-conditioned place preference in these male offspring. This work demonstrates for the first time that microglia play a key role in translating prenatal opioid exposure to long-term changes in neural systems and behavior.

**Highlights:**

- Prenatal opioid exposure decreases offspring viability and body weight in males and females
- Prenatal opioid exposure decreases microglial phagocytosis of D1R in the nucleus accumbens in males only
- Prenatal opioid exposure increases nucleus accumbens dopamine D1 receptor expression in males but not females
- Adult males fail to extinguish oxycodone-conditioned place preference following prenatal oxycodone exposure

## Introduction

In the past decade, rates of opioid use disorder have increased to epidemic proportions (Haight et al., 2018). This includes a dramatic increase in disorders among pregnant mothers, and therefore the incidence of prenatal exposure in their children. Indeed, the Centers for Disease Control and Prevention report that between 1999 and 2014 the incidence of opioid use disorder at delivery increased by 333% (Haight et al., 2018). Infants exposed to opioids during gestation often develop neonatal opioid withdrawal syndrome (NOWS) which is characterized by tremors, difficulty feeding, high-pitched crying, inconsolability, and diarrhea (Conradt et al., 2019; Arter et al., 2021). While these symptoms can be acutely managed with opioid replacement therapies, very little is known about the long-term consequences of prenatal opioid exposure for brain development and behavior.

The existing literature on the long-term effects of prenatal opioid exposure, while sparse, points towards enduring consequences. For example, differences have been found between school age children exposed to opioids in utero and un-exposed children in intelligence quotient (IQ)/cognitive performance (Nygaard et al., 2015; van Baar et al., 1994), visual acuity (Moe, 2002), language (Hunt et al., 2008; Benninger et al., 2020), attention-deficit disorder and/or hyperactivity (Sandtorv et al., 2018), and aggression and anxiety (de Cubas & Field, 1993). Nygaard et al. (2017) found that 17–to-21-year-olds prenatally exposed to opioids scored significantly lower on a number of cognitive performance metrics as compared to un-exposed peers. Importantly, however, interpretation of the human literature is confounded by several factors, including retrospective chart review, small sample sizes, and variables such as maternal nutrition, access to health care, socio-economic status, and maternal mental health (Singer et al., 2020; Conradt et al., 2019). Interestingly, Nygaard et al. (2015), using a prospective study design, found that boys, but not girls, exposed to opioids in utero had lower IQ scores at 8 years of age. Similarly, boys, but not girls, have poorer language and cognitive scores following prenatal opioid exposure during early childhood (Skumlien et al., 2020). These findings suggest the potential for a male bias in susceptibility to the long-term effects of prenatal opioid exposure.

A wide array of behavioral and neurobiological changes have also been identified in animal models of prenatal opioid exposure (Grecco et al., 2021; Minakova et al., 2021; for review see Byrnes & Vassoler, 2018; Abu & Roy, 2021, Boggess & Risher, 2020). For example, adult male rats exposed to morphine during gestation exhibit decreased extinction of methamphetamine-conditioned place preference (CPP), as well as increased drug-primed reinstatement of CPP, as compared to controls (Shen et al., 2016). Similarly, adolescent male rats demonstrated increased behavioral sensitization to methamphetamine following prenatal methadone (Wong et al., 2014). The mesolimbic reward circuit is a key neural substrate on which opioids act to induce changes in behavior. In the nucleus accumbens (NAc), a key target region of dopaminergic inputs from the ventral tegmental area (VTA), endogenous opioids and dopamine facilitate the hedonic and motivational aspects of drug seeking behavior, respectively. The effects of prenatal opioid exposure on the endogenous opioid system have been studied extensively (for review see Byrnes & Vassoler, 2018), but much less is known about how prenatal opioid exposure influences the development of the dopamine system within the NAc. Both dopamine D1 and D2 receptor (D1R and D2R) densities in the NAc peak during adolescence (at approximately postnatal day [P]30) in male rats, and then decline to adult levels by P55 (Kopec et al., 2018; Tarazi & Baldessarini, 2000). Microglia, the resident immune cells of the brain, are increasingly recognized to play a key role in the developmental sculpting of neural circuits. We recently showed that microglial pruning of D1Rs is responsible for this developmental decline following the peak at P30, but only in males (Kopec et al., 2018). Early work has begun to explore the impact of prenatal opioid exposure on microglia. Prenatal methadone exposure reduces the ramification state of microglia in the cortex at P10 (often taken as an indicator of increased proinflammatory activation) and increases brain expression of toll-like receptor 4 (Tlr4) and its adaptor molecule myeloid differentiation primary response protein 88 (MyD88; Jantzie BBI 2020). Furthermore, prenatal exposure to methadone increased serum concentrations of the proinflammatory cytokines interleukin 1ß (IL-1ß), tumor necrosis factor alpha (TNFα), interleukin 6 (Il-6), and the chemokine CXCL1 at P10, and the increase in IL-1ß persisted until P21 (Jantzie BBI 2020).

Based on these collective findings, we hypothesized that prenatal opioid exposure would impact microglial function and disrupt D1R pruning, leading to persistent alterations in the NAc dopamine system, potentially in a sex-specific manner. To test our hypothesis, we used intravenous oxycodone self-administration in rats to approximate a real-world scenario in which women are chronically self-administering opioids – due either to an untreated opioid use disorder or to medication-assisted treatment (e.g., methadone, buprenorphine) for an opioid use disorder. Female rats self-administered the prescription opioid, oxycodone, for 3 weeks prior to mating to account for the fact that women do not initiate opioid use upon pregnancy, but rather become pregnant in the context of their opioid use. We then had female rats continue to self-administer oxycodone throughout pregnancy until parturition, when access to drug was stopped.

## Methods

### Animals

Adult male and female Sprague-Dawley rats (Charles River Laboratory, Wilmington, MA) were group-housed under standard laboratory conditions (12-h light-dark cycle [lights on at 7:00am], food and water ad libitum). All experiments were approved by and conducted in accordance with the Animal Care and Use Committee at McLean Hospital and the National Institutes of Health guide for the care and use of Laboratory animals.

### Intravenous oxycodone self-administration

Adult female rats underwent jugular catheter implantation surgery and were trained to self-administer oxycodone hydrochloride (NIDA Drug Supply) prior to mating. Briefly, rats were implanted with chronic, indwelling silastic (0.51mm internal diameter) intravenous jugular catheters as described in Mavrikaki et al., 2017. Catheters were cleaned daily by flushing with 0.2mL of heparinized saline and disinfected once/week with 0.2mL gentamicin (10mg/ml). One week after surgery, females underwent oxycodone self-administration training in operant conditioning chambers (Med Associates, St. Albans, VT, 30.5 × 24.1 × 29.2cm). Operant chambers were enclosed in sound-attenuated cubicles with ventilation fans and contained 2 retractable response levers with cue lights, a house light, a counterbalanced fluid swivel and tether, and an infusion pump. A syringe containing oxycodone solution (0.1mg/kg) was located outside the chamber and connected to the rat’s catheter via a spring-covered Tygon tube connected to a fluid swivel. Rats self-administered oxycodone for 4-h/d on a Fixed Ratio 1 (FR1) schedule of reinforcement, with one press on the active lever resulting in a 4-second infusion (0.1ml/infusion), the concentration of which was adjusted to the rat’s body weight every 3 days. Infusions were followed by a 6-second time out period in which rats could press the levers, but no drug infusion occurred. These doses and procedures are adapted from (Vassoler & Byrnes, 2019). Prior to mating, females were given access to self administration for 4hrs/day, 5 days/week for 3 weeks. Control dams were implanted with jugular catheters and allowed to self-administer oxycodone for 5 days in the first week, after which they remained in their home cages with no more access to oxycodone. Next, both the control and oxycodone-exposed females were time-mated, and pregnancy was determined by confirmation of sperm in the vaginal canal (assessed by microscopic evaluation of vaginal swabs). Following mating, the oxycodone treatment group continued to self-administer oxycodone 7-days/week until the day before parturition. Control dams remained in their home cages. Oxycodone self-administration was halted upon parturition.

### Neonatal pup outcomes and behavioral quantification

Number of pups in each litter was counted on P1, P3, and P6 to assess neonatal mortality. In addition, milk band size was assessed in all pups between 9:00am-10:00am on P1 as a measure of how much milk they were receiving from the dams during the night and categorized according to a 1-4 rating scale (1: no band, 2: small band, 3: medium band, 4: large band). Data are presented as % in each rating category per litter. Body weight was also assessed in all offspring at P1, P3, P6 and then again from P21-P50 (P21, P26, P36, P46, P50) to determine long-term changes in weight gain. Neonatal behavioral tests were conducted to assess motor capacities (righting reflex and incline plane). The righting reflex test was conducted at P1 and P3. In this test, pups were placed on their backs on a flat surface, and the latency for each pup to right itself was recorded with a stopwatch. Latency to right and the number of pups and percent per litter unable to right were quantified. The incline plane test was conducted at P6. In this test, pups were placed facing downward on a 45° angle plane and the latency for each pup to pivot 180° so they were facing upwards was recorded with a stopwatch. Pups were scored based on the final angle reached (1: 0-90°, 2: 90-179°, 3: 180°) as well on latency to reach 180° (if attained). Both the righting reflex and incline tests had a maximum cut-off time of 60 seconds. Data for male and female pups were combined for these measurements, as our ability to accurately sex the pups on P1-P6 was not reliable.

### Brain Tissue Collection & Sectioning

At P20, P30, and P50, both male and female offspring from both control and oxycodone dam groups were anesthetized using a mixture of ketamine/xylazine and transcardially perfused using ice-cold saline followed by 4% Paraformaldehyde (PFA). Brains were postfixed in 4% PFA for 48 hours followed by 30% sucrose for 48 hours and then frozen at -80°C before cryosectioning at 40µm on a cryostat. Brain sections were stored in cryoprotectant (sucrose, polyvinylphrrolidone, and ethylene glycol) at -20°C until IHC staining. Ages for assessment were chosen based on our previous work (Kopec et al., 2018) to span the adolescent period and to capture the peak timepoint for microglial pruning of D1R (P30).

### Immunohistochemistry (IHC)

IHC staining was conducted according to Kopec et al. (2018). Briefly, every 5^th^ section within the NAc was chosen for any given stain (200µm apart; Bregma: +2.76mm to +2.28mm). Brain sections were rinsed 5 times in 1xPBS and then incubated at 80°C for 30 min in 10mM sodium citrate (pH 9.0) for epitope retrieval. Next, sections were incubated in 1mg/ml sodium tetraborate in 0.1M PB and then 50% methanol in PBS to quench background fluorescence for 1 hour each. Sections were blocked for 1 hour in 10% goat normal serum with 0.3% Triton-x100 and 3% H202 in 1x PBS before primary antibody incubation. Two separate IHCs were conducted in separate NAc brain sections to label for D1R + microglia (using Iba1) and for dopamine D2 receptor (D2R) and tyrosine hydroxylase (Th; labels dopamine fibers). For D1R and Iba1 staining, sections were incubated for 48 hours at 4°C with a mouse D1R antibody (Novus Biologicals #NB110–60017; Littleton, CO; 1:1500) and a chicken Iba1 antibody (Synaptic Systems, 1:1500). We previously validated the specificity of the Novus D1R antibody in our own hands using D1R knock-out mouse tissue (Kopec et al., 2018). These were followed by secondary antibody incubation with goat anti-mouse Alexa-Fluor (AF)488 and goat anti-chicken AF568, respectively (Thermofisher scientific, each 1:500) for 2 hours at room temperature (RT). For D2R and Th staining, sections were incubated for 24 hours at RT with a rabbit D2R antibody (Milllipore, 1:1500) and a mouse Th antibody (Immunostar, 1:1000). These were followed by secondary antibody incubation with goat anti-rabbit AF568 and goat anti-mouse AF488, respectively (both Thermofisher Scientific 1:500). Following IHC, all sections were mounted on gelatin subbed slides, coverslipped with vectashield antifade mounting media with DAPI (Vector Labs), sealed with nail polish, and stored at -20°C until imaging.

### Imaging and analysis of D1R, D2R, and Th immunofluorescence

To quantify immunofluorescent intensity in the NAc, 40x magnification Z-stacks were taken on a Zeiss AxioImager microscope. 10 step Z-stacks were taken with a step-size of 0.65µm measured from the center of the focal Z-plane. Images were taken medial to the anterior commissure (AC) using the AC as a landmark. A total of 3-6 images were taken between Bregma +2.76 and +2.28 for each animal (based on the Paxinos and Watson Rat Brain Atlas). For D1R, D2R, and Th, fluorescence intensity in each image was analyzed using FIJI. Briefly, Z-stacks were converted to maximum projection images and mean grey value was measured and normalized to background (defined as the mean grey value of the AC) using the entire images as the RO1. The values of all images taken for a given animal were averaged to provide a single measurement per rat at each age.

### IMARIS quantification of microglial engulfment of D1R and microglial morphology

To quantify microglial engulfment of D1R at P30, 63X magnification Z-stacks were taken on a Zeiss Airyscan 880 Confocal Laser scanning microscope. Z-stack step size was 0.3 µm and each image had 60-100 steps, aiming to capture entire microglia. Imaris 9.5.1 (Bitplane Scientific Software) was used to create surface renderings of individual microglia (Iba1 labeling) and D1R (D1R labeling) within the microglial surface. Volume of engulfed D1R was then quantified and normalized to total cell volume. 4-5 cells were reconstructed per animal from 3-5 separate brain sections between Bregma distances of +2.76 and +2.28 within the NAc. For assessment of microglial morphology, IMARIS neurite tracer was used to create a skeletonization of individual microglia on which Sholl analyses were conducted.

### Place conditioning with oxycodone in adult offspring

To begin to test whether prenatal exposure to oxycodone affected reward function in adult offspring, we used place conditioning to asses the sensitivity of adult male and female offspring to oxycodone-induced conditioned place preferences. Briefly, an unbiased, three-compartment place-conditioning apparatus (Med Associates) was used according to Russell et al., 2016. Within the apparati, each chamber differed in lighting (bright vs. dim), floor texture (mesh vs. metal rods), and wall coloring (black vs. white). On the first day, a “screening” session was conducted in which rats were placed into the apparatus and given 20 min to freely exlore all three compartments. Rats that showed any inherent bias (i.e. preference for one of the compartments ≥ 14 min.) were eliminated from further testing. On the second day, rats underwent two place-conditioning sessions. In the morning (10:00am) conditioning session, all experimental rats were injected with saline and confined to one side compartment within the apparatus for 30 min, counter balanced across sides and groups. In the afternoon (2:00pm) conditioning session, all experimental rats were injected with oxycodone (3mg/kg) and confined to the opposite side compartment for 30 min. On the third day, experimental rats were given 20 min to freely explore all 3 compartments in a single “test” session. This was followed by 2 days of additional test sessions (extinction sessions 1 and 2) in which experimental rats were exposed to the apparatus without any treatments and allowed to freely explore for 20 min.

### Statistics

Graphpad Prism version 9.2.1 was used for all statistical analyses. A one-way ANOVA with repeated measures on time was used to assess oxycodone self-administration in dams. 2-way mixed-effects ANOVAs (age x treatment) were used to assess number of pups/litter, avg. body weight/litter, body weight per animal, and righting reflex outcomes (% of litter unable to right and avg. latency to right/litter). Milk band size (%/litter) and inclined plane outcomes (avg./litter) were compared using un-paired t-tests. For D1R and Th mean grey values, 2-way ANOVAs (age x treatment) were used, followed by Bonferroni post-hoc tests, in each sex separately because tissues from males and females were processed separately and thus, could be compared directly. For D2R mean grey values at P55, t-tests were used to compare treatment groups. For microglial engulfment analyses, nested t-tests were used to compare treatment groups. For Sholl analyses, nested t-tests were used. 2-way mixed effects ANOVAs (session x treatment) were used to assess conditioned place preference. All data are represented as mean +/- SEM and significance was set at p<0.05.

## Results

### Prenatal oxycodone exposure impacts neonatal outcomes in offspring

Experimental dams self-administered progressively more oxycodone across pregnancy (F_(35,499)_=1.66, p=0.01, Fig. 1A). In offspring, prenatal opioid exposure significantly decreased the number of surviving pups per litter at P3 and P6 (F_(1, 18)_= 8.66, p<0.01, Fig. 1B, posthoc P3: p<0.01, P6: p<0.01). This is potentially due to impaired nursing, as ∼25% of oxycodone pups had no visible milk band, and there was a trend towards a smaller proportion of pups with milk bands classified as medium sized (t_(1,18)_=1.961, p=0.066, Fig. 1C). Average pup body weight per litter was also significantly lower during the perinatal period (F_(1,18)_=7.276, p<0.05, Fig. 1D), and this decrease persisted into adulthood (F_(3,131)_=67.63, p<0.0001, Fig. 1E). Importantly, this persistent decrease in body weight was observed in both males and females (posthoc males: p<0.05, females p<0.001).

**Figure 1.**
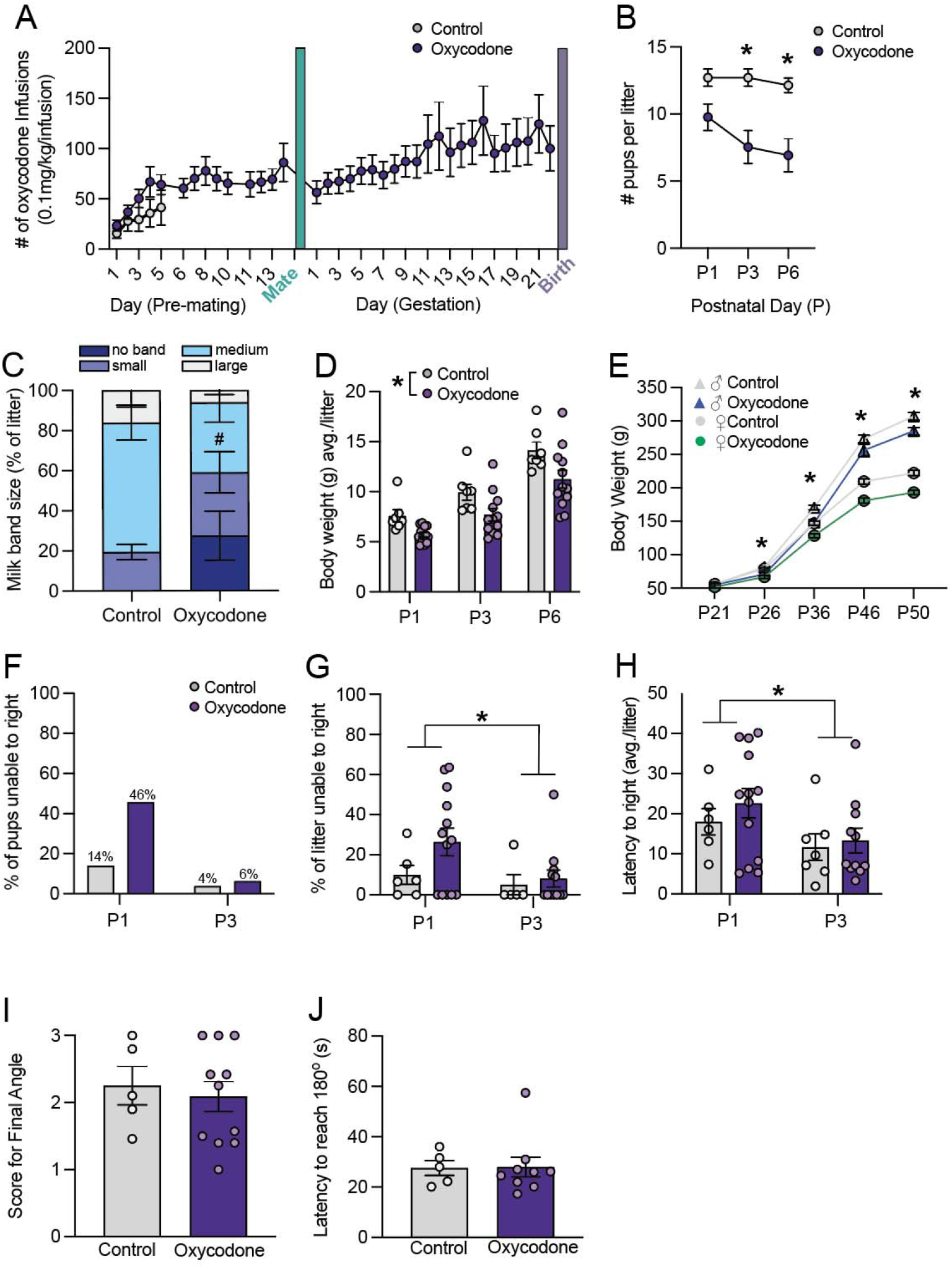
Prenatal opioid exposure impacts neonatal outcomes in offspring. **A)** Rat dams self administered oxycodone for either 5 days pre-pregnancy (Controls) or throughout gestation. **B)** Number of pups per litter was significantly decreased at P3 and P6 following oxycodone exposure. **C)** ∼ 20% of oxyocodone exposed pups lacked a detectable milk band and oxycodone pups tended to have fewer medium-size milk bands. **D)** We observed a main effect of treatment such that the average body weight of oxycodone exposed litters was smaller than that of control litters. **E)** Both male and female oxycodone exposed offspring were persistently lower in body weight into adulthood. A-D: N = 7 litters (CON), 12 litters (Oxycodone), male and female offspring are combined. **F)** 46% of oxycodone exposed pups were unable to right themselves in 60 sec. on P1, as compared to only 14% of control pups, N=79-127 pups per group. **G)** There was no significant treatment effect on the % of each litter unable to right or on the latency to right **(H)**, N (litters) = 6 (P1 control), 5 (P3 control), 13 (P1 oxycodone), 12 (P3 oxycodone). However, both measures were significantly lower at P3 as compared to P1. **I)** There was no difference between control and oxycodone pups in the final angle reached in the inclined plane test, or in the latency to reach 180° **(J)**. Data represent Mean +/- SEM, *p=0.05.

The righting reflex and inclined plane tests were conducted at P1 and P3 or P6, respectively. We observed only mild effects of prenatal oxycodone treatment on these behavioral outcomes. 45% of oxycodone pups were unable to right themselves within 60 sec. at P1, as compared to 14% of control pups – suggesting motor impairments (Fig. 1F). However, the percentage of pups in each litter that were unable to right in 60 seconds only trended towards significance at P1 (t_(1,17)_=1.528, p=0.14; Fig. 1G). The average latency to right (per litter) was higher at P1 as compared to P3 (Age effect: F_(1,33)_=4.302, p<0.05, Fig. 1H) but did not differ between treatment groups (Treatment effect: F_(1,33)_=0.689, p=0.413, Fig. 1H). In the inclined plane test, there was no effect of oxycodone on either the final angle reached on average/litter (t_(1,14)_=0.428, p=0.675, Fig. 1I) or on the average latency to reach 180° (t_(1,12)_=0.2064, p=0.95,Fig. 1J).

### Prenatal oxycodone exposure increases D1R density in the NAc in adulthood in males but not females

Our previous work showed that D1R density is higher in the NAc at P30 as compared to younger (P20) or older (P55) timepoints in male rats (Kopec et al., 2018). We hypothesized that prenatal oxycodone exposure might alter the developmental trajectory of D1R in the NAc between P30 and P55.Thus, we assessed D1R immunofluoresent intensity in the NAc at P20, 30, and 55 (Fig. 2A & B). We found a significant effect of age (p<0.05) and a significant age x treatment interaction effect (p<0.05) for D1R (See Table 1 for complete statistics). Posthoc testing revealed a significant increase in D1R at P55 in oxycodone treated males relative to controls (Fig. 2C&E, p<0.05). Importantly, this increase was male-specific as there was a main effect of age for D1R in the females (Fig. 2F&H, p<0.001), but no treatment or interaction effects. At P55, there was no difference in D2R between control and oxycodone exposed males (Fig. 2D, t_(11)_=0.563; p=0.585) or females (Fig. 2G, t_(11)_=0.536; p=0.602), demonstrating that D1R but not D2R, is increased in the NAc by prenatal opioid exposure. To determine whether changes in D1R were driven by changes in dopaminergic input to the NAc, we also assessed the density of Th fibers within the NAc at the same timepoints (Fig. 2i). We found significant effects of age in both males (Fig. 2J, p<0.01) and females (Fig. 2K, p<0.001), but no treatment or interaction effects. These findings led to our hypothesis that decreased microglial phagocytosis of D1R, and not changes in DA fiber input to the NAc, is responsible for higher NAc-D1R following prenatal oxycodone exposure.

**Table 1.** Complete results of 2-way ANOVAs (treatment x age) for D1R and Th in male and female offspring with Bonferroni posthoc tests. Values represent p values, bolded if significant at p<0.05. Color coding indicates significant age comparisons (blue) and treatment comparisons (green).

| Comparison: | D1R Males | Th Males | D1R Females | Th Females |
| --- | --- | --- | --- | --- |
| Main effect: Treatment | **F (1, 30) = 9.766, p=0.004** | F (1, 29) = 0.752, p=0.752 | F (1, 32) = 0.309, p=0.582 | F (1, 32) = 0.379, p=0.542 |
| Main effect: Age | **F (2, 30) = 3.983, p=0.029** | **F (2, 29) = 6.500, p=0.005** | **F (2, 32) = 10.83, p<0.001** | **F (2, 32) = 9.090, p<0.001** |
| Interaction Effect: Treatment x Age | F (2, 30) = 2.265, p=0.121 | F (2, 29) = 0.941, p=0.402 | F (2, 32) = 2.166, p=0.131 | F (2, 32) = 0.680, p=0.514 |
| Posthoc PD20 vs PD30 | **p=0.031** | **p=0.041** | **p<0.001** | **p=0.001** |
| Posthoc PD30 vs PD55 | p>0.999 | **p=0.006** | **p<0.001** | **p=0.007** |
| Posthoc PD20 vs PD55 | p=0.155 | p>0.999 | p>0.999 | p>0.999 |
| Posthoc Control vs. Oxycodone PD20 | p>0.999 | p>0.999 | p=0.963 | p>0.999 |
| Posthoc Control vs. Oxycodone PD30 | p=0.897 | p>0.999 | p=0.201 | p=0.676 |
| Posthoc Control vs. Oxycodone PD55 | **p=0.003** | p=0.675 | p>0.999 | p>0.999 |

**Figure 2.**
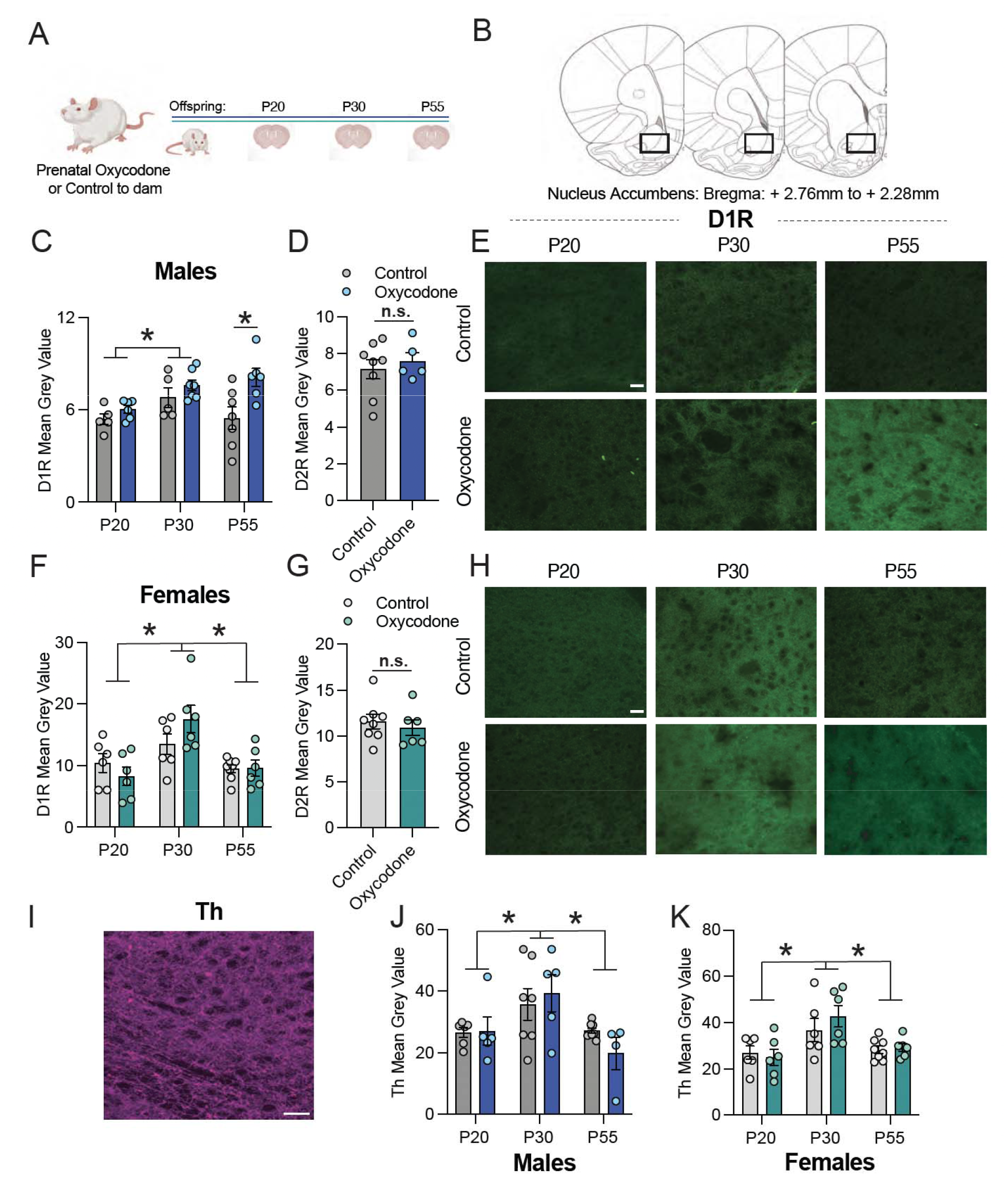
Prenatal oxycodone exposure increases D1R density in the NAc in adulthood in males but not females. **A)** Schematic of timeline for treatment and tissue collection. **B)** Rat brain atlas images delineating NAc region where D1R, D2R, and Th were quantified (based on Paxinos and Watson Rat Brain Atlas). **C)** In males, D1R mean grey value differed significantly with age, and was higher at P55 in oxycodone exposed males as compared to control. **D)** D2R did not differ with treatment at P55 in males. **E)** Representative 20x images of D1R staining in the NAc of males, scale bar= 27 microns. **F)** D1R peaked at P30 in both control and oxycodone exposed females, but did not differ with treatment. **G)** D2R did not differ with treatment at P55 in females. **H)** Representative 20x images of D1R staining in the NAc of females, scale bar= 27 microns. **I)** Representative 20 image of Th staining in the NAc of males, scale bar = 27 microns. **J)** Th peaked at P30 in both control and oxycodone exposed males and females **(K)**, but did not differ with treatment. Data represent Mean +/- SEM, *p=0.05.

### Prenatal opioid exposure decreases microglial engulfment of NAc-D1R during adolescence in males

We previously showed that microglial complement-dependent phagocytosis of D1R is required for the normal developmental decline in D1R between P30 and P55 in male rats. Moreover, we found that local pharmacological blockade of microglial phagocytosis – specifically at P30 – within the NAc, increased D1R at PD55 (Kopec et al., 2018). Therefore, we hypothesized that higher NAc-D1R following prenatal oxycodone exposure might be due to decreased microglial phagocytosis of D1R at P30 (Fig. 3A). In line with this hypothesis, we found that D1R volume within microglia at P30 was significantly lower following prenatal oxycodone exposure (Fig. 3B&C, % D1R volume/microglia, t_(11)_=2.36; p<0.05), while microglia volume did not differ between treatment conditions (Fig. 3D&E, t_(11)_=0.97; p=0.36). Interestingly, this effect did not appear to be due to gross changes in microglial morphology, as Sholl analysis revealed no significant differences in number of branch endpoints in microglia (Fig. 3F&G, t_(11)_=0.96; p=0.39) or in Schoenen’s Ramification Index (Fig. 3H, t_(11)_=0.68; p=0.51).

**Figure 3.**
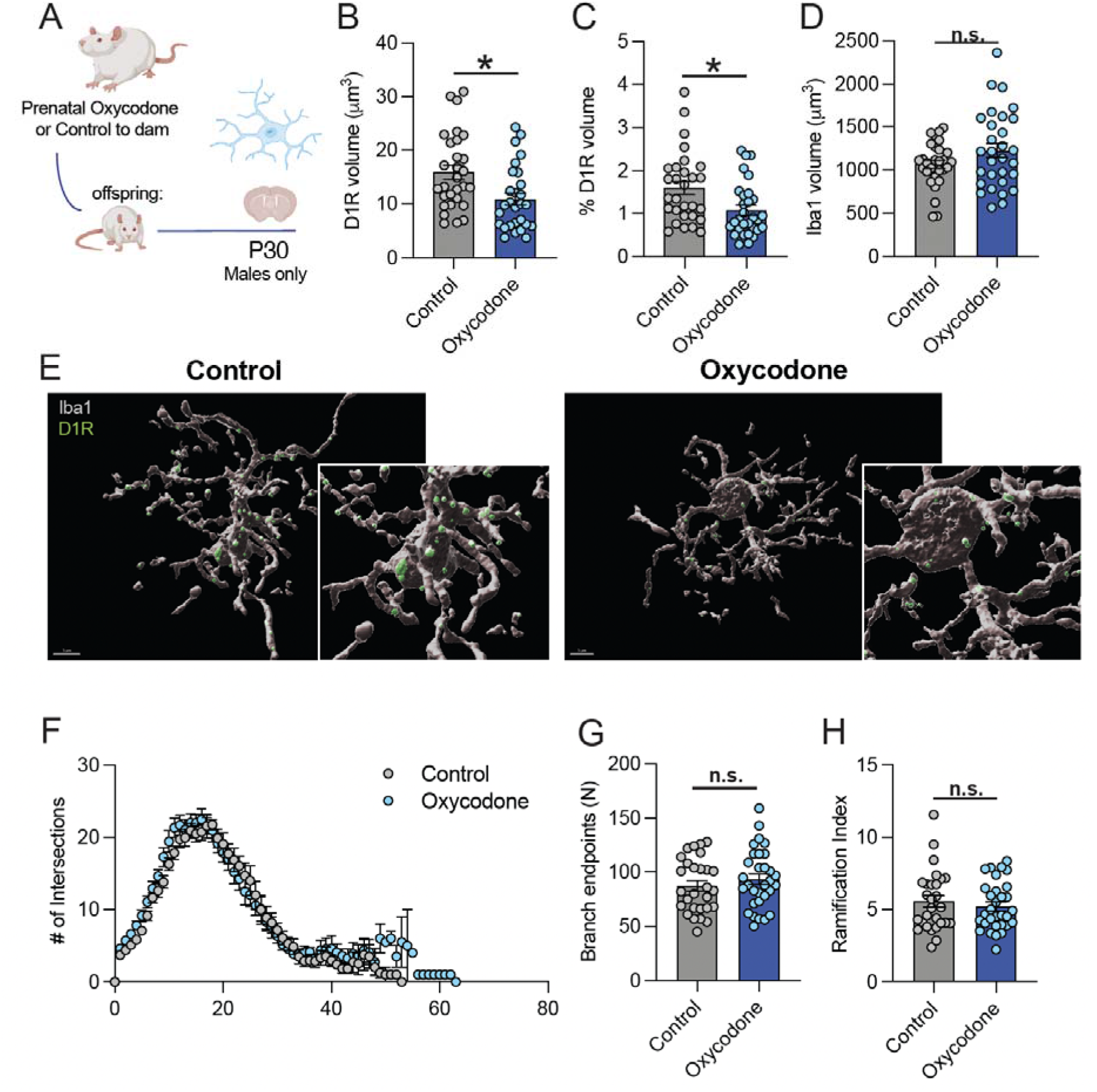
Prenatal opioid exposure decreases microglial engulfment of NAc-D1R during adolescence in males. **A)** Schematic of timeline for exposures and P30 engulfment assessment. **B)** D1R volume within microglia was significantly decreased following oxycodone exposure in male offspring at P30. **C)** % D1R volume of total microglial volume was significantly decreased following oxycodone exposure in male offspring at P30. **D)** There was no treatment effect on microglial volume. **E)** Representative 60X images of microglial D1R engulment: grey: Iba1, green: D1R, scale bar= 5 microns. **F)** Scholl analysis revealed no differences in # of intersections between control and oxycodone males, or in branch endpoints **(G)** or Ramification Index **(H)**. B-H: dots represent individual microglia, N=29 (control), 30 (oxycodone) nested per animal 6 and 7, respectively for analysis (nested t-tests). Data represent Mean +/- SEM, *p=0.05.

### Prenatal opioid exposure reduces extinction, but not acquisition, of oxycodone-induced conditioned place preferences in adult male but not female offspring

In order to begin to determine lasting behavioral effects of prenatal opioid exposure on adult behavior, we assessed both the acquisition and extinction of oxycodone-induced conditioned place preferences in male and female offspring (Fig. 4A). In males, we found that a significant main effect of session (F_(2,22)_=12.24, p<0.01, Fig. 4B) as well as significant session x treatment interaction effect (F_(2,22)_=3.79), p<0.05), but no significant main effect of treatment (F_(1,11)_=0.30, p=0.596). Posthoc testing revealed that while expression of oxycodone-induced conditioned place preferences decreased significantly between test and extinction day 1 in control males (p<0.01), oxycodone-exposed males failed to extinguish their preference for the oxycodone-paired compartment – as evidenced by no significant differences between test day and either extinction day 1 or 2. In females, there was a significant main effect of session (F_(2,20)_=11.88, p<0.01, Fig. 4C), but no significant main effect of treatment (F_(1,10)_=0.030, p=0.865) or session x treatment interaction effect (F_(2,20)_=2.271, p=0.129) indicating that both control and oxycodone-exposed female offspring demonstrated normal extinction of oxycodone-induced conditioned place preferences.

**Figure 4.**
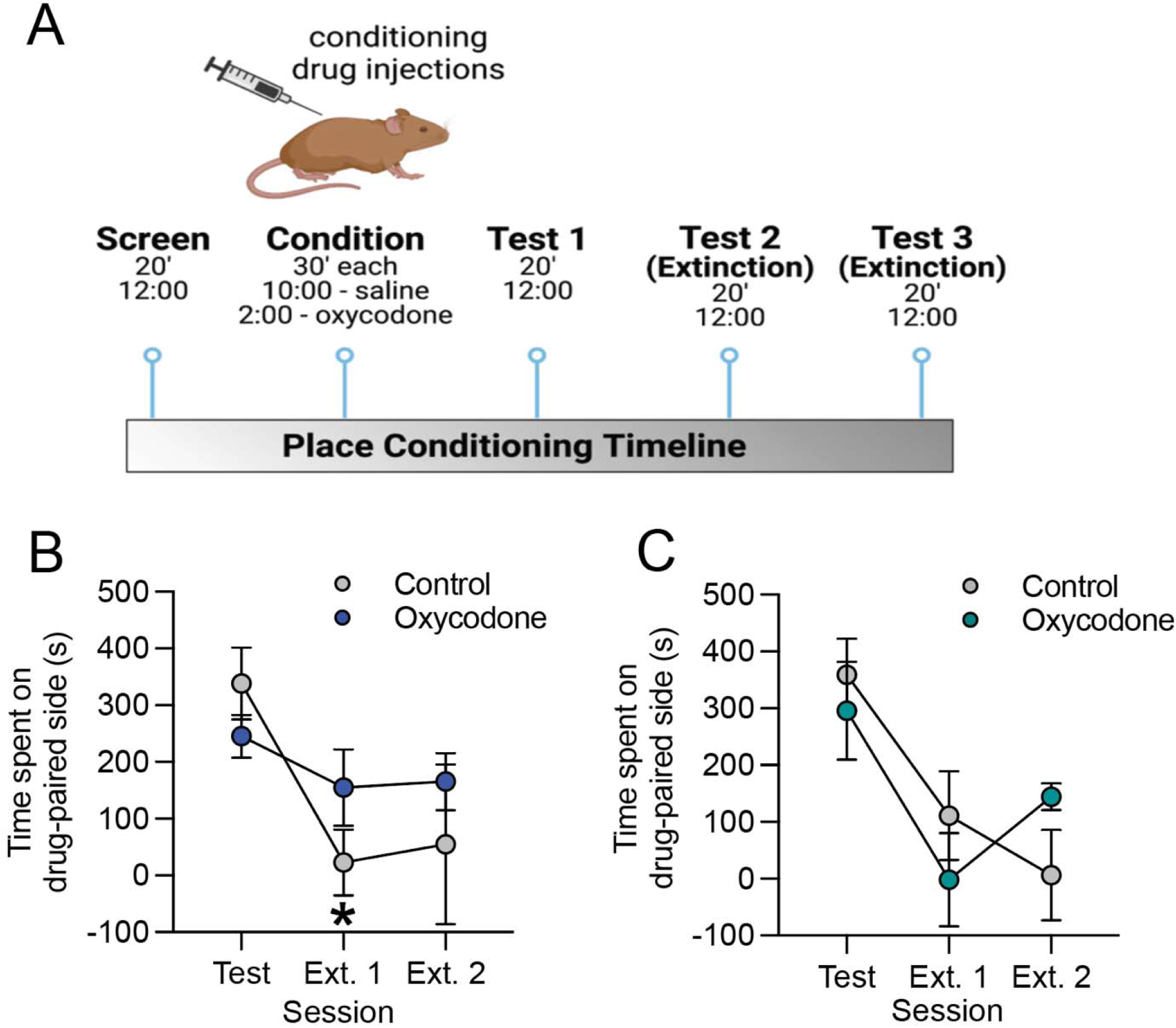
Prenatal opioid exposure prevents extinction of oxycodone-conditioned place preference in adult male, but not female, offspring. **A)** Schematic of conditioned place preference procedure. **B)** Control male offspring showed a significantly lower preference for the drug-paired compartment on extinction session 1 as compared to the test session, while no decrease between test and extinction 1 was observed in oxycodone-exposed males. **C)** Both control and oxycodone-exposed females showed extinction of conditioned place preference. Data represent Mean +/- SEM, *p=0.05 relative to Test session.

## Discussion

Our results demonstrate that prenatal oxycodone exposure increases D1R density in the NAc of young adult male rats (P55) compared to controls. Notably, this effect is sex-specific as no increase was observed in females. Neither D2R density nor Th-immunoreactive (ir) fiber density in the NAc differed between control and prenatal oxycodone exposed groups, suggesting that higher D1R density is not driven by greater Th inputs to the NAc. Rather, microglial engulfment of D1R is reduced at P30 following prenatal oxycodone exposure in males – a timepoint which we have previously determined is a critical age at which microglia phagocytose NAc-D1Rs (Kopec et al., 2018). Together, these findings suggest that prenatal opioid exposure impairs the ability of microglia to prune D1R during adolescence in males. Consequently, the normal developmental decline in NAc-D1R between adolescence and adulthood fails to occur, resulting in increased D1R density in males only.

Microglia are increasingly recognized to play a critical role in the developmental sculpting of neural circuits in the brain, by phagocytosing synaptic elements and even whole newborn neurons (Faust et al., 2021; Vanryzin et al., 2019). Moreover, human studies have implicated microglia as a key cell type in substance use disorders. For example, RNAsequencing of postmortem brain tissues from individuals with a history of substance use revealed changes in inflammatory gene sets, including TNFα signaling pathways (an important proinflammatory pathway in microglia) specifically within the NAc (Seney et al., 2021). However, virtually nothing is known about how prenatal exposure to drugs of abuse alters microglial phagocytic processes. To our knowledge, our findings are the first demonstration that prenatal opioid exposure disrupts a normal microglial developmental neural circuit pruning program. Indeed, very little work has been done to investigate the impact of prenatal opioid exposure on microglia or neuroimmune function in general. Jantzie et al., (2020) found that perinatal methadone exposure (from embryonic day 16 to P21) increased serum proinflammatory cytokines at P10 including IL-1ß, IL-6, TNFα, and CXCL1. Importantly, this increase in circulating IL-1ß persisted to P21. They also reported that TLR4 and MyD88 mRNA were elevated in brain issues from offspring at P10, as were IL-1ß and CXCL1 protein. Finally, they found that cortical microglia were rounder with less branch complexity in methadone-exposed offspring as compared to controls at P10 (Jantzie et al., 2020). One important caveat to this work is that the methadone exposure was mostly during the postnatal period, so it is unclear how this is comparable to the prenatal period. For instance, in contrast to Jantzie et al., we did not observe changes in microglial morphology following prenatal opioid exposure. However, it is entirely possible that such a difference could be due to factors such as brain region, type of opioid, and/or age at exposure. Further work is needed to fully characterize the impact of prenatal opioid exposure on microglial function and the relevance of such changes to neural circuit function and behavior.

An outstanding question is how prenatal opioid exposure alters microglial pruning during adolescence. Studies have investigated the molecular pathways by which opioids impact microglial biology in adolescent and adult animals or cell cultures. In adolescent rats, the expression of the chemokines CCL4 and CCL17, as well as their receptor CCR4, are upregulated in microglia isolated from the NAc, following morphine exposure (Schwarz et al., 2013). Moreover, rats that receive repeated morphine during adolescence (but not young adulthood) show persistent changes in microglial function and increased reinstatement to morphine CPP as adults; and, pre-treatment with a glial modulator during adolescent morphine exposure prevents this increased reinstatement, implicating a critical role for microglia (Schwarz et al., 2013), but the precise mechanisms were not defined. In adult male mice, TLR2 is upregulated in the brain by morphine treatment, and TLR2 knock-out (KO) mice have reduced microglial activation in response to morphine (Zhang et al., 2011). In adult male mice, Rivera et al. (2019) found that microglia-specific KO of MyD88 increased microglial phagocytosis of newborn (doublecortin positive) neurons in the hippocampus following morphine exposure. Rivera et al. (2019) also showed that male mice lacking microglial-MyD88 showed prolonged extinction of morphine-conditioned place preference and increased drug reinstatement. Our results herein, along with those of Shen et al. (2016), demonstrate that prenatal exposure to opioids decreases extinction of either oxycodone- or methamphetamine-conditioned place preference (respectively) in adulthood. Together, these findings suggest that microglia are an essential mediator between prenatal opioid exposure and the reward-related effects of drugs of abuse in adulthood.

While few other studies have investigated the impact of opioids on microglial phagocytic processes, insight can be gained from literature on peripheral tissue-resident macrophages which share many similarities with microglia. In humans, opioids have been shown to induce immunosuppression in opioid users and, therefore, increased vulnerability to infections, in part by inhibiting macrophage phagocytosis of bacteria (Ninkovic et al., 2016; Kozlowski et al., 2019; Rojavin et al., 1993). Tomassini et al. (2003) found that this opioid inhibition of macrophage phagocytosis appears to be mediated via mu and delta opioid receptor signaling (Tomassini et al., 2003). Similarly, mu opioid knock-out mice do not display opioid-induced reductions in macrophage phagocytosis (Roy et al., 1998). Based on these findings, it is interesting to speculate that mu-opioid receptor signaling might represent a potential mechanism by which down-regulation of microglial D1R phagocytosis is initiated. It will also be important to investigate the epigenetic programs by which prenatal opioid exposure exerts effects on later-life microglia behavior.

In conclusion, our results provide novel evidence that prenatal opioid exposure disrupts the development of the dopamine system, at least in part by altering microglial engulfment of D1R during adolescence. Importantly, this is only the case in males; no changes in the dopamine system parameters that we assessed were altered by prenatal opioid exposure in females. Further work is needed to shed light on the molecular mechanisms by which these changes occur, as well as their long-term implications for behavior. Finally, the factors that provide resilience in the face of opioid exposure in females remain to be elucidated.

## Acknowledgements

We would like to thank all the members of the Bilbo lab for their critical reading of the manuscript, as well as the Animal Care staff at McLean Hospital for providing excellent animal care and the staff of the Duke Light Microscopy Core for assistance with learning and trouble-shooting confocal microscopy.

## Author contributions

E.H.C., S.D.B., C.J.S., and T.L. designed the study. C.J.S., T.L., M.J.C., K.E.M., N.C., A.A., Y.A.C., V. J.K., and Y.C.J. conducted experiments. C.J.S, E.H.C., and S.D.B. wrote the manuscript. All authors approved of the final version of the manuscript.

